# Fokker-Planck diffusion maps of multiple single cell microglial transcriptomes reveals radial differentiation into substates associated with Alzheimer’s pathology

**DOI:** 10.1101/2024.06.21.599924

**Authors:** Andrew Baumgartner, Max Robinson, Todd Golde, Suman Jaydev, Sui Huang, Jennifer Hadlock, Cory Funk

## Abstract

The identification of microglia subtypes is important for understanding the role of innate immunity in neurodegenerative diseases. Current methods of unsupervised cell type identification assume a small noise-to-signal ratio of transcriptome measurements that would produce well-separated cell clusters. However, identification of subtypes is obscured by gene expression noise, diminishing the distances in transcriptome space between distinct cell types and blurring boundaries. Here we use Fokker-Planck (FP) diffusion maps to model cellular differentiation as a stochastic process whereby cells settle into local minima, corresponding to cell subtypes, in a potential landscape constructed from transcriptome data using a nearest neighbor graph approach. By applying critical transition fields, we identify individual cells on the verge of transitioning between subtypes, revealing microglial cells in inactivated, homeostatic state before radially transitioning into various specialized subtypes. Specifically, we show that cells from Alzheimer’s disease patients are enriched in a microglia subtype associated to antigen presentation and T-cell recruitment.

## Introduction

High throughput single-cell RNA sequencing (scRNA-seq) technologies and analysis techniques have enabled the profiling at single-cell resolution of gene expression of individual microglia cells in patients with Alzheimer’s disease. Olah et. al. [1] measured the transcriptome of 16,000 microglia cells from post-mortem and fresh brain samples in patients with AD, mild cognitive impairment (MCI), and temporal lobe epilepsy (TLE), identifying 9 microglia sub-types, with some reportedly depleted in AD patients. More recently, Patel et al. also reported 9 microglial subtypes, profiling over 26,000 cells from 5 donors and multiple pathologies, including AD [2]. Mathys et al. performed single cell transcriptomics on mixed brain cell types, reporting another microglial subtype depleted in AD patients with similar, but distinct, transcriptional profiles [3]. Prater et. al. [4] recently uncovered 10 microglia transcriptomics substates using single-nucleus RNAseq, some of which were altered in Alzheimer’s disease. Together, these studies have shown that transcriptional profiling of microglia can help uncover homeostatic and altered substates, and identify specific genes and pathways unique to these substates that can potentially be targeted for future treatments for AD. These reports represent a stream of analyses that extend the initial studies on microglia transcriptional profiling in both human and mice [5], by offering improved resolution of microglia subtypes. Here we present methods that capture the dynamics of the generation of the subtypes to further refine the resolution to identify substates and that improve robustness of discrimination.

While the aforementioned studies have provided a substantial leap forward in understanding microglial subtypes, they do not provide a picture of their differentiation dynamics, i.e. how these subtypes are developmentally related to each other.Although subtypes can provide valuable insights into mechanisms of disease, there is no “ground truth” by which to assess the validity of the proposed subtypes. Standard scRNAseq analysis has proved useful in uncovering the transcriptional differences between cell types, but little work has been done in assessing the biological validity of hence identified subtypes. Existing workflows often involve unsupervised clustering of cell transcriptomes followed by marker gene identification, and thus, make the implicit assumption that the magnitude of fluctuations of gene expression profiles are much smaller than the distance in transcriptome space between clusters, and that the characteristic time scale of these fluctuations are much faster than that for differentiation. These assumptions stem from the underlying dynamical system view that during differentiation cells relax into well-defined minima in some “cellular phase space”. These minima would, in loose analogy, reflect some “low energy potential” or high-probability wells [6]. However, bulk transcriptome data of sorted outlier cells at the border of clusters) have shown that fluctuations slow enough [7] for these two time scales to merge, creating, non-reproducible cluster boundaries that lack proper mathematical justification. Put simply, there’s no clear way to draw the lines between subtypes.

The characteristic lengths (in transcriptome space) and times associated to such cell state dynamics, including the distances between the minima and the relaxation times, provide set of scales by which we can reliably assess the effects of intrinsic and extrinsic gene expression noise on the resulting cell (sub)type classification. If the magnitude of fluctuations is small in comparison to the distance between minima, it is unlikely that transitions between these minima will be due to noise alone. This will be reflected in the data by well separated clusters. On the other hand, if the noise is comparable to the difference between the minima, then stochastic fluctuations will frequently drive cells between the minima, thereby flattening out the underlying potential landscape giving rise to ill-separated clusters [8]. This small noise approximation is well justified if there are large transcriptional differences between cell types, as in inter-cell type scRNA-seq studies, with well separated clusters. However, such differences are parametrically smaller when studying intra-cell type transcriptional subtypes, or substates, and the small noise approximation may not be appropriate, leading to misclassification of individual cells, arbitrary distinctions between clusters, and formation of spurious clusters.

In this work, we set out to examine the dynamics of microglia substates by finding a low dimensional representation of the underlying dynamical system via Fokker-Planck diffusion maps [9, 10, 11, 12, 13] and interpreting the results in the continuum limit. We apply our approach to an integrated set of single-cell transcriptome data by Olah and Patel [1, 2]. We chose an integrated dataset to both increase the number of cells, as well as reducing potential bias from a single dataset. This integration of data from two sources increases the number of cells and reduces the potential of bias. A thorough theoretical treatment of FP diffusion maps, with illustrations and applications to single cell transcriptome data can be found in [13]. This approach allows us to reclassify the substates directly in terms of the underlying dynamical systems, and to identify long lived, robust cellular states. The validity of these states comes from the ability of diffusion maps to separate long-time dynamics from noise in systems which admit a spectral gap. Differential expression analysis and interpretable machine learning on these substates point to specific reactive responses of microglia to putative AD pathology and display qualitative behaviors similar to previously reported microglia substates. Additionally, the structure of the low dimensional representation combined with repurposed tools developed for longitudinal bulk RNAseq implies that these substates differentiate radially from a central, inactive, homeostatic state.

## Results

Before presenting our results, we would like to make explicit the assumptions underlying the validity of our results. The primary, and perhaps most tenuous, assumption is that cells can be modeled as continuous vectors of normalized gene expression values. That is, cells live in a continuous vector space. There has been interesting recent work in modeling stochastic single cell RNA expression via master equations [14, 15] which aims to challenge this assumption. While we appreciate this rigorous work, we know there exist a limit in which the master equations give rise to a set of deterministic differential equations where this assumption holds true. Such a limit is known as the “system size expansion” [16], where the precise definition of system size depends on the problem at hand. For chemical reaction networks, such as those underlying biological gene expression, this system size is traditionally the volume of the solution in which the reactions are taking place. However, in single cell analysis, we are primarily concerned with modeling the *relative* counts of genes in a given cell [17]. This implies that the natural system size for stochastic transcriptomics is the total UMIs in a given cell: *N* . In [18] it was shown that fluctuations about the deterministic dynamics (i.e when fluctuations are small) are of order 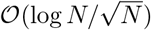. The same work shows that the system including the first order fluctuations can be modeled by a diffusion process up to 𝒪 (log *N/N* ). This is by no means a perfect approximation, but it is acceptable so long as we keep cells with total UMIs∼ 𝒪 (10^3^), or the number of cells with UMI < 10^3^ is much smaller than the number of cells with UMIs > 10^3^. This gives fluctuations around 10% to 30% of the mean values for genes of 𝒪(10). Thus our continuum assumption primarily captures the dynamics associated to the most highly expressed genes in our cell population. Additionally, the diffusion approximation requires the timescale of differentiation to be much shorter than the typical lifetime of a microglia cell, allowing us to treat the number of microglia as constant and ensuring the validity of our Fokker-Planck analysis. The responsiveness of microglia to external stimuli and the evidence that the number of microglia remain relatively constant throughout one’s lifetime [19] suggest that this assumption is a valid one. Additionally, the diffusion approximation requires each cell be in local equilibrium with respect to the Waddington landscape. This assumption is related to the existence of a gap in the spectrum of the graph Laplacian, whereby degrees of freedom below the gap equilibrate, leaving only the long-time dynamics.

### Fokker-Planck diffusion map reveals radial structure to microglia substate dynamics

Using single cell mRNA counts for microglia from Olah et al. and Patel et al. [1, 2] integrated using Seurat v4 [20], we created Fokker-Planck (FP) diffusion maps [13] to observe the differentiation dynamics across the population of cells. Diffusion maps consider the underlying physics that positions individual cells in some cellular phase space, such as gene expression space. In brief, the single cell transcriptomic states are determined for each cell by the abundance of all the *M* mRNAs species in a cell that collectively form the cell’s gene expression vector *x* of *M* components. The state *x* of a cell thus maps to a point in an *M* -dimensional gene expression state space at a position that uniquely represents the cell’s transcriptome. Since each cell is represented by a point in said space a change of the transcriptome in a cell is a movement in gene expression state space. The gene expression noise that randomly alters the gene expression profile is then manifest as random wiggling of individual cells, which spreads a set of nominally same cell types into a cloud or cluster. Interpreting the dispersion of cells in transcriptome space, and related “distances” that they travel a consequence of gene expression noise underlies the idea diffusion maps as an alternative to the UMAP dimension reduction that is devoid of a physical basis.

In contrast to noise, gene expression profiles of cells can also change in a concerted manner, due to regulatory signals that cause cell differentiation, which shifts gene expression profile in a net direction, towards that which represent the new cell type – in state space this is manifest as a net movement of a group of cells. Such deterministic herding of cells to a distinct new position, called drift, happens on top of the stochastic diffusion; and both the deterministic drift in a given direction and the dispersion by noise are jointly modelled by the Fokker-Plank (FP) equation. We have recently proposed an implementation of diffusion maps that explicitly models these processes based on stochastic models of transcriptional dynamics [13] using a discretization of the (backward) Kolmogorov equation. Figure 1 shows both a UMAP [21] annotated by our cluster labels and several views of the first four diffusion coordinates.

**Figure 1.**
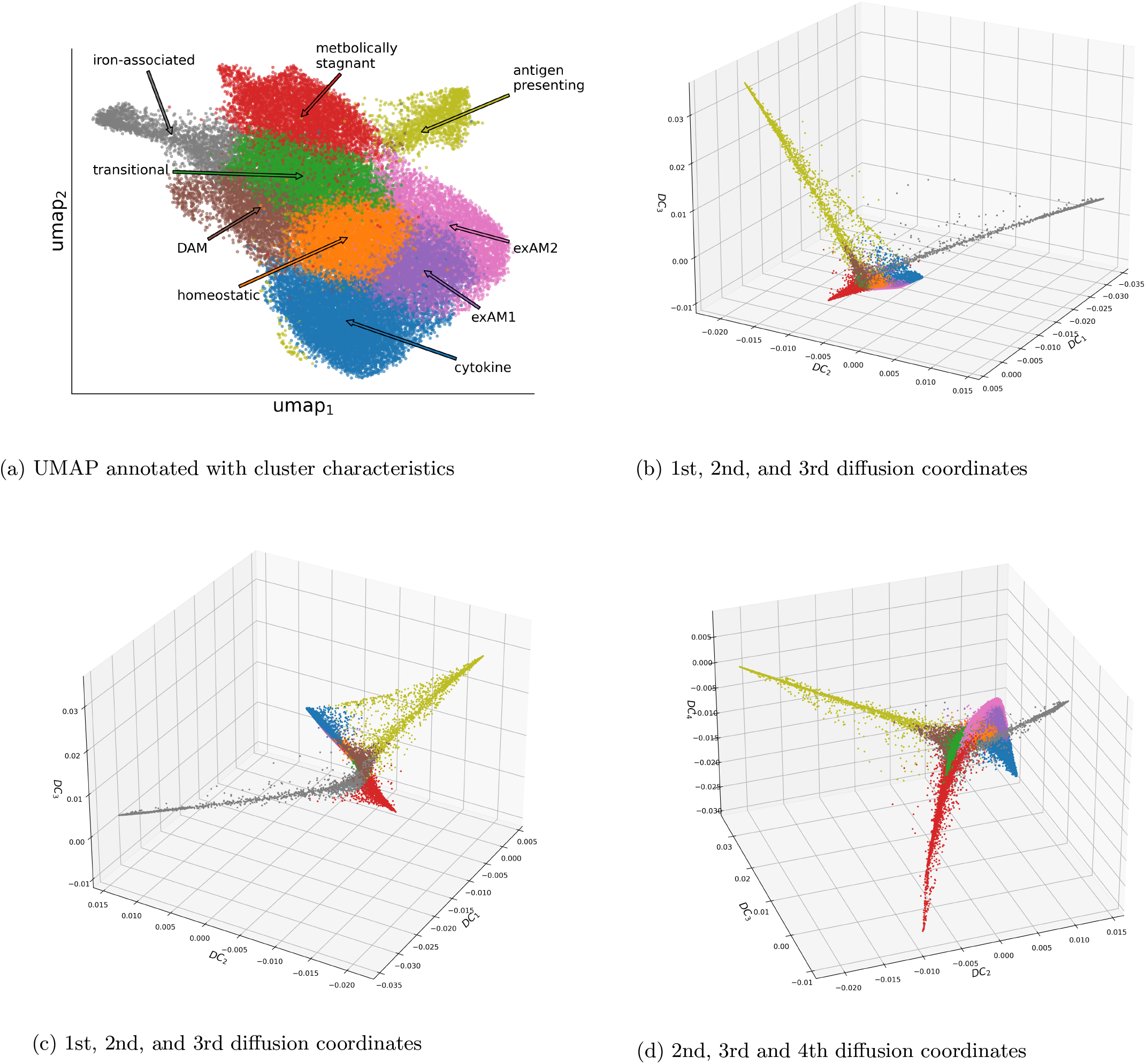
UMAP and Fokker-Planck diffusion maps colored by microglia subtype found using Leiden clustering. The prominent clusters visible in the first few diffusion coordinates are the largest minima and therefore the most robust to noise induced perturbation (See Methods). The UMAP is strictly for visualization purposes.

More formally, the FP diffusion map has at its core the following elementary process in which the gene expression state of individual cells evolves in time according to the following stochastic differential equation

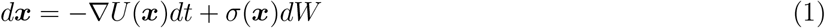

Herein, *x* represents the (normalized) gene expression vector of individual cells. The first term −∇*U* (*x*)*dt* captures aforementioned drift that causes net move of cells, and the second term represents the gene expression noise that disperses the cells. Because the detailed functional form of the drift is not known (it would have to come from the gene regulatory pathway) yet, cell differentiation appear phenomenologically to follow some sort of a landscape that afford an imaginary force unto cells, best illustrated by Waddington’s epigenetic landscape. Thus, a commonly accepted and successfully applied approximation for the drift is to view it as driven by a gradient on a slope of a landscape [22, 23, 24, 25, 26] whose topography captures biological phenomenology and is the equivalent of a free energy or potential landscape that assigns a scalar value *U* (*x*) to each position, or gene expression state *x*. The gradient function ∇ turns this scalar quantity at *x* into a vector pointing to the steepest direction in the landscape as if driven by gravity, – the change of transcriptome to which cells are driven to as preordained by the gene regulatory network. The second term *σ*(*x*)*dW* represents gene expression noise – the source of the diffusion component, with *σ*(*x*) being the state-dependent strength of the noise, and *dW* representing Wiener (Gaussian) noise.

Using our proposed approach (see Methods and [13]) one can read off the shape of this landscape directly from the single-cell transcriptome data. For cell populations, we are interested the probability distribution of cells in gene expression state space, whose individual behavior is governed by eq. (1), as function of time and how they can form the observed distribution manifest as clusters in the data. For this purpose we use the backward Kolmogorov equation associated to eq. (1), the adjoint of the Fokker-Planck equation on the space of all *L*^2^ integrable functions. To fit the observed gene expression data to the Kolmogorov equation we use techniques described in the [13] for uncovering the landscape geometry through appropriately connecting all the cells to their nearest neighbors and diffusion maps.

### Critical transition fields and pseudo-radial ordering indicate radial differentiation dynamics

The dynamical representation of all the cells in various states as a snapshot in the FP diffusion maps in fig. 1 shows an apparent radial structure with a central cluster surrounded by robust substates extending along each of the diffusion coordinate, suggesting that the latter arise from radial diversification of the central cluster. While this interpretation based on the structure in the FP diffusion map is appealing and, to some extent, warranted this structure alone, derived solely from a snapshot data, is not enough to establish an actual radial differentiation pattern emanating from the central cluster. To do this we rely on the fact that in the deterministic (zero noise or large *N* ) limit, transitions into stable attractor states occur via a bifurcation in the underlying dynamical system [23]. A simple way to determine if a bifurcation is close to occurring from snapshots of single-cell transcriptomes was laid out in [27, 28] and along similar lines of reasoning in [29, 30], taking advantage of increasing dispersion of cells when they leave destabilized potential minima, while aligning the gene expression values in *x*. Given a data matrix ***X*** where *X*_*ij*_ is value of the *j*^*th*^ gene in the *i*^*th*^ cell the critical transition parameter is

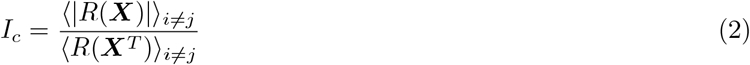

where *R*(·) is the matrix of correlation coefficients of the input matrix and ⟨·⟩_*i*≠*j*_ is the average over all off-diagonal elements. It is shown in [27] that the numerator vanishes and the denominator diverges when the system is in a local minima while the opposite behavior occurs during a bifurcation. When one performs a linear transformation such as projecting onto the principal components, this divergence is rendered finite but the qualitative behavior remains. We generalize this parameter into a field over cells by calculating the correlation coefficients in a small neighborhood about each cell. That is, the averages in eq. (2) are performed among ***x***’s *k* nearest neighbors and smoothed using a median convolution filter. We find that cells with large values of Ic are located primarily in our proposed central cluster (Fig. 2), indicating that it is the source of diverging differentiation into the more robust subtypes and thus, has a “homeostatic” function as source to replenish the various specialised cell populations. In our FP diffusion map, these cells are present in high density in a small ball around the origin.

**Figure 2.**
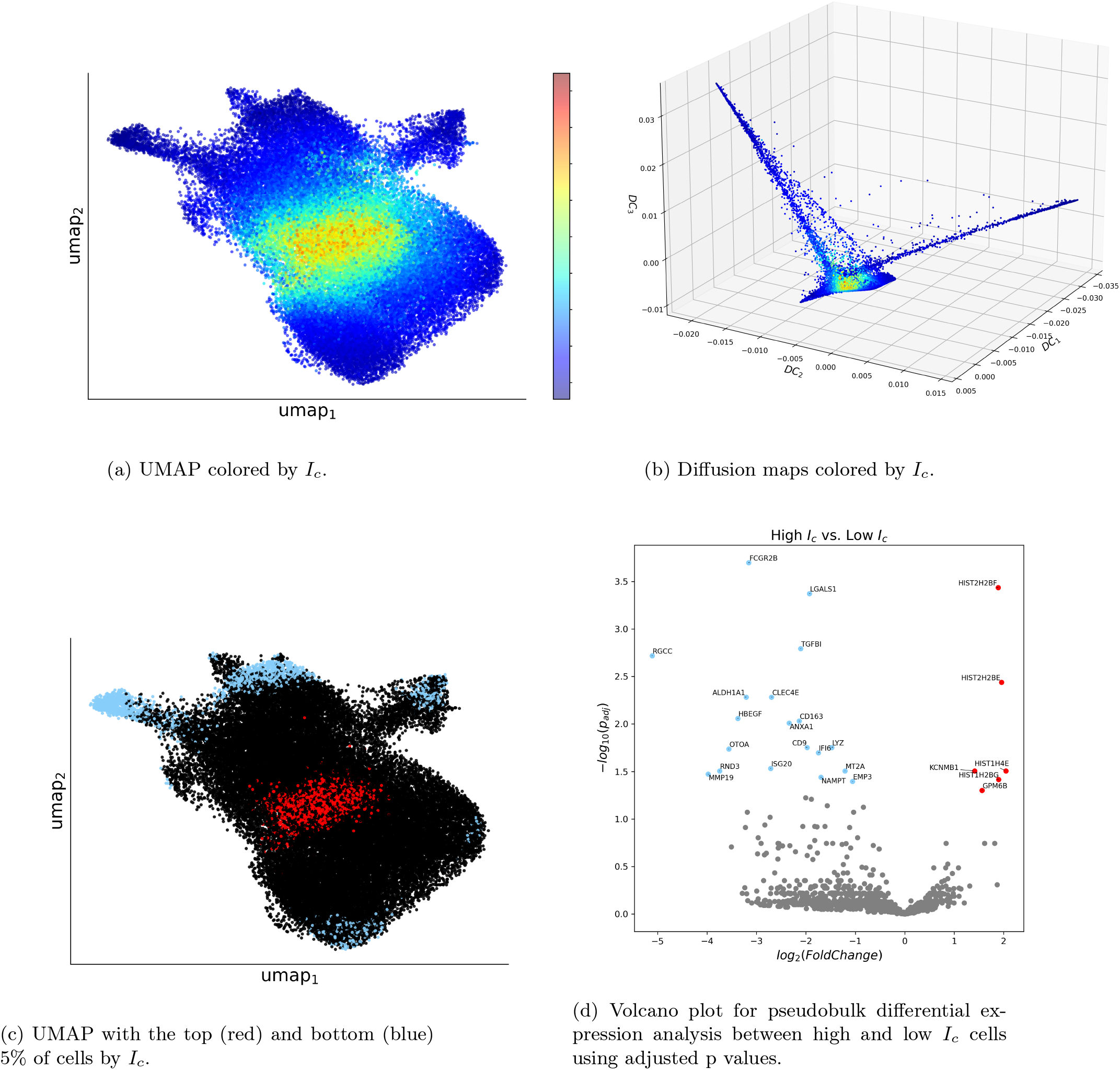
UMAP and diffusion map colored by *I*_*c*_. Large values of *I*_*c*_ correspond to cells which are about to undergo and cell state transition, while small values correspond to cells which are in or near an attractor state.

The concentration of large *I*_*c*_ cells in the center of the FP diffusion map is again suggestive of a radial differentiation pattern. To corroborate this interpretation, we use diffusion pseudotime [31] to impose a pseudo-*radial* ordering on the data with respect to the centroid of the diffusion map projection and plot *I*_*c*_ as a function of this ordering for each cluster. As figure 3 shows, cells with large values of the critical transition displayed small pseudoradius and those with small values had large pseudoradius, indicating further away along a respective axis of differentiation and state stabilization.

**Figure 3.**
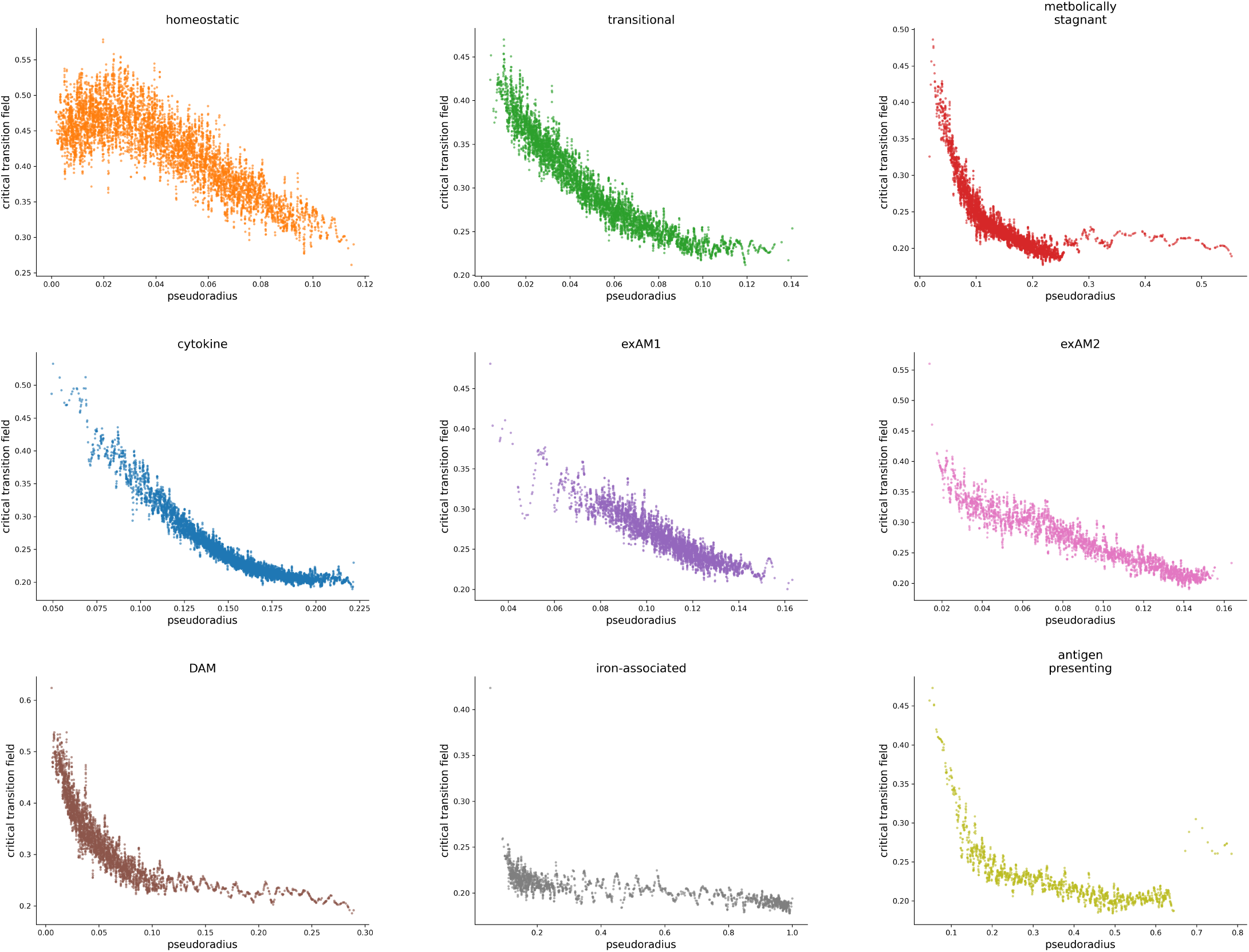
Ic values in each cluster as a function of the pseudoradius smoothed using a Savitzky-Golay filter. The pseudoradius is measured with respect to the centroid of the FP diffusion maps, defined as the point in the first three diffusion coordinates with the largest local density estimate (see Methods). Decreasing values as a function of the pseudoradius implies cell along these trajectories are undergoing state transitions and settling into their specialized microglia subtypes.

**Figure 4.**
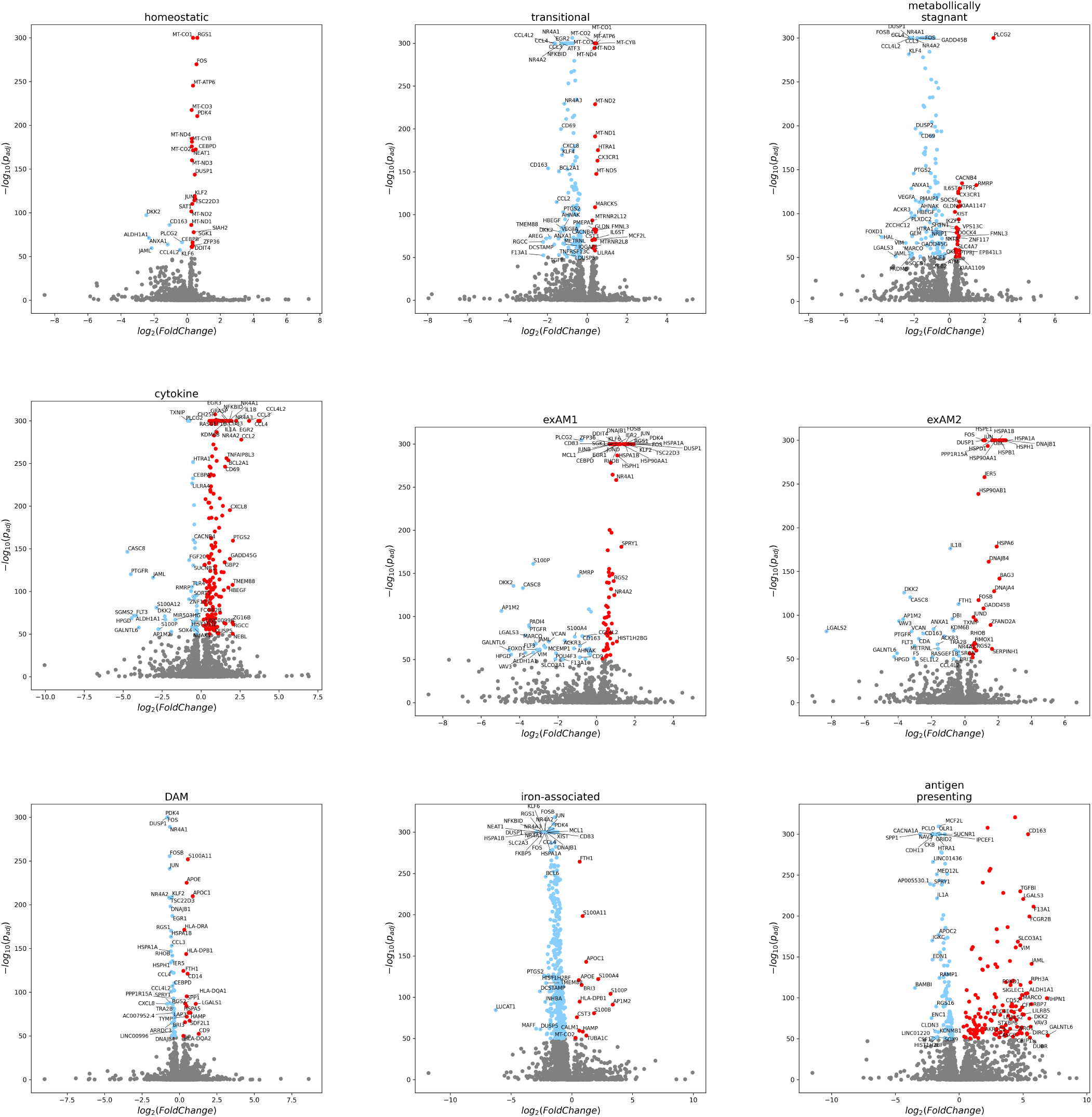
Volcano plots for one vs. all single-cell differential expression analysis. The top 50 up and down regulated genes by log fold change with *p*_*adj*_ < 10^−50^ are annotated.

Finally, to expose the specific transcriptional changes as *I*_*c*_ decreases, we performed pseudobulk differential expression analysis between the top and bottom 5% of cells based on *I*_*c*_. By using pseudobulk profiles of the mixture of cells from each subtype, we identified global changes common to all transitions out of the homeostatsic state (Fig. 3d) Interestingly, several members of the histone H2B1 and H2B2 families (HIST1H2BG, HIST2H2BE, HIST2H2BF) were upregulated in large Ic cells, as well as a member of the histone H41 family (HIST1H4E). While the core histones H2B transcripts mark S phase, thus proliferating cells – consistent with their role as source progenitor cells, non-core histone isoforms play a role in chromatin accessibility regulation, notably in conjunction with epigenetic cell fate control. Interestingly, TGFBI, a protein that inhibits cell adhesion, was significantly downregulated, consistent with the notion that microglia are more mobile in their functional state.

### Differential expression and interpretable machine learning reveal subtype functions

To identify characteristic genes in each cluster we performed a “1 vs. all” analysis using scanpy’s default method [32]. We found a very high degree of overlap in differentially expressed genes in each cluster, consistent with the common theme in tissues that cell subtypes are not marked by exclusive subtype-specific genes but the the pattern of relative expression of genes. This can be seen in figure 7 where even after z-scores are calculate, there is significant heterogeneity for all genes across subtypes.

While differential expression reveals the molecular characteristics of a given subtype, it does not identify the genes that act as discriminatory feature of the subtypes. To identify such genes we built a gradient boosted tree classifier (XGBoost) to classify the single cells into our subtypes using the top differentially expressed genes as features. The confusion matrix for our model is displayed in Fig. 5 a) while the ROC curves for this one vs. rest classification is shown in 5 b). We achieved balanced accuracy of 81.5% and a combined ROC-AUC score of about .98. We calculated feature importance using SHAP values [33] for each cluster to determine the most important genes for classification into that cluster(Fig. 6). Note that the high or low feature values referred to in the SHAP plots is about the z-scored values of the gene within that cell. High feature values thus represent high z-scores, not the absolute expression of the gene, but rather it’s expression relative to all the other differentially expressed genes.

**Figure 5.**
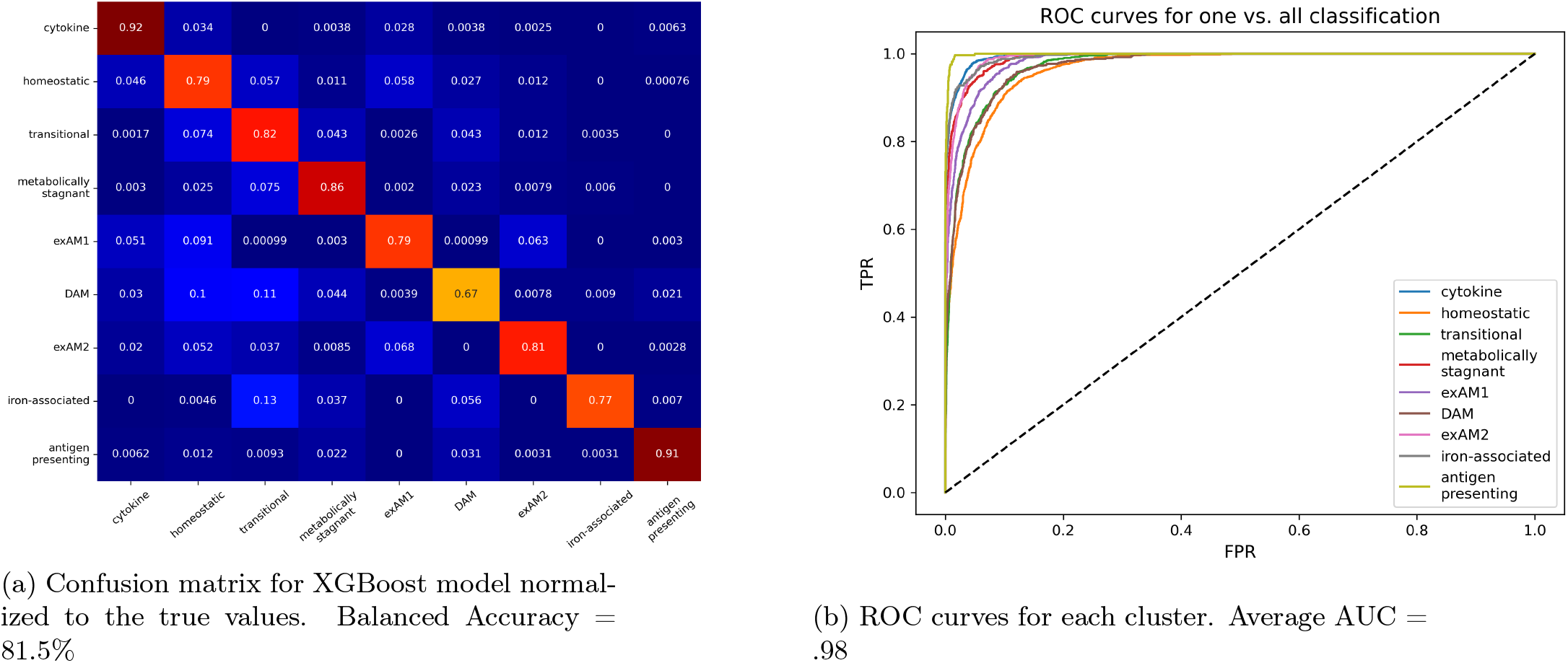
Performance metrics for our XGBoost model for classifying cells into their respective suybtypes. These models were trained on z-scored values of signatures genes displayed in figure 7. The criteria for these genes are that they must be deferentially expressed in at least 50% of cell in the given subtype.

**Figure 6.**
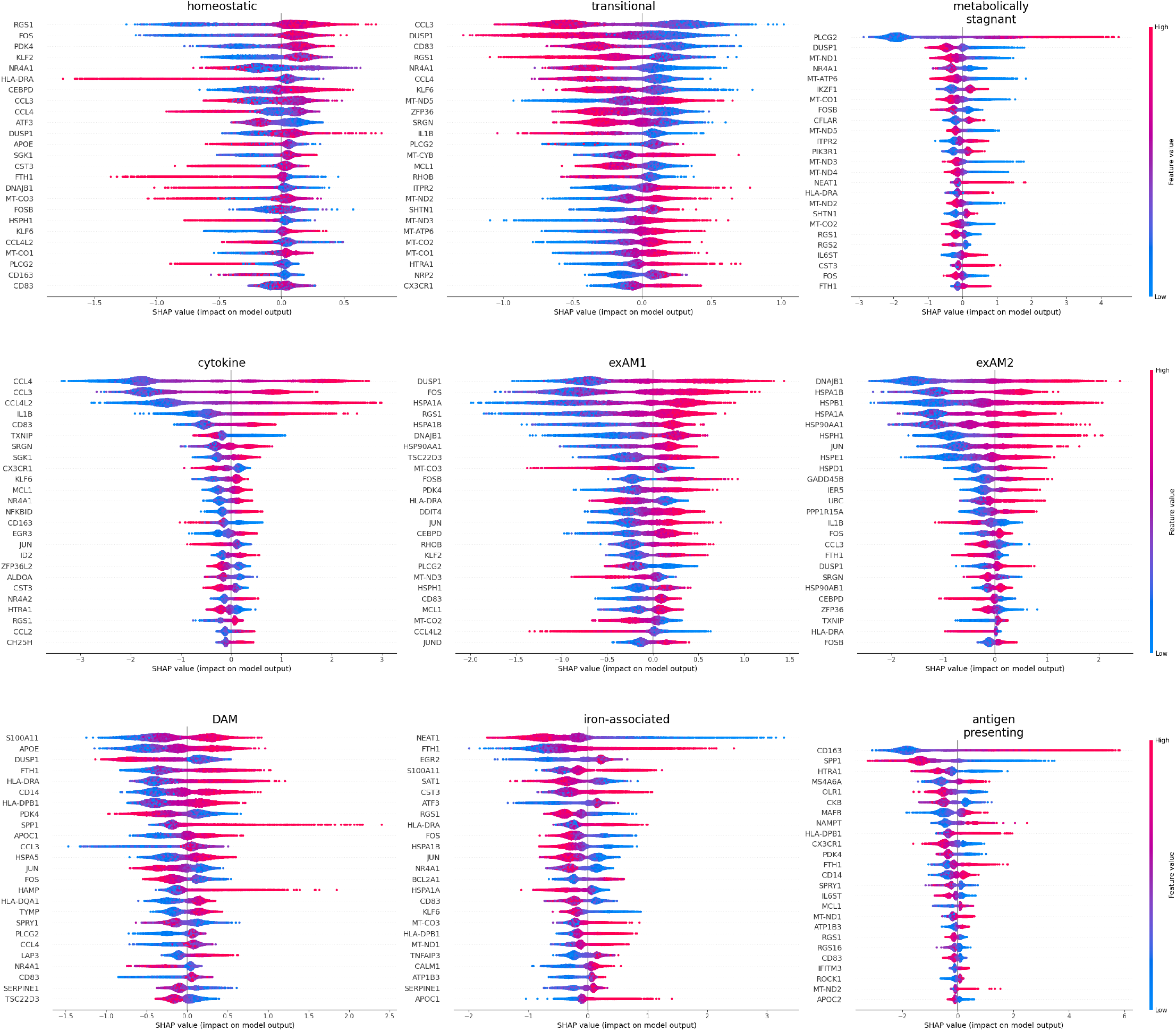
SHAP summary plots for each cluster. Genes with positive SHAP values with high features values indicates high expression of the gene relative to the other signtures genes, and are important for classifiying cells into that cluster.

**Figure 7.**
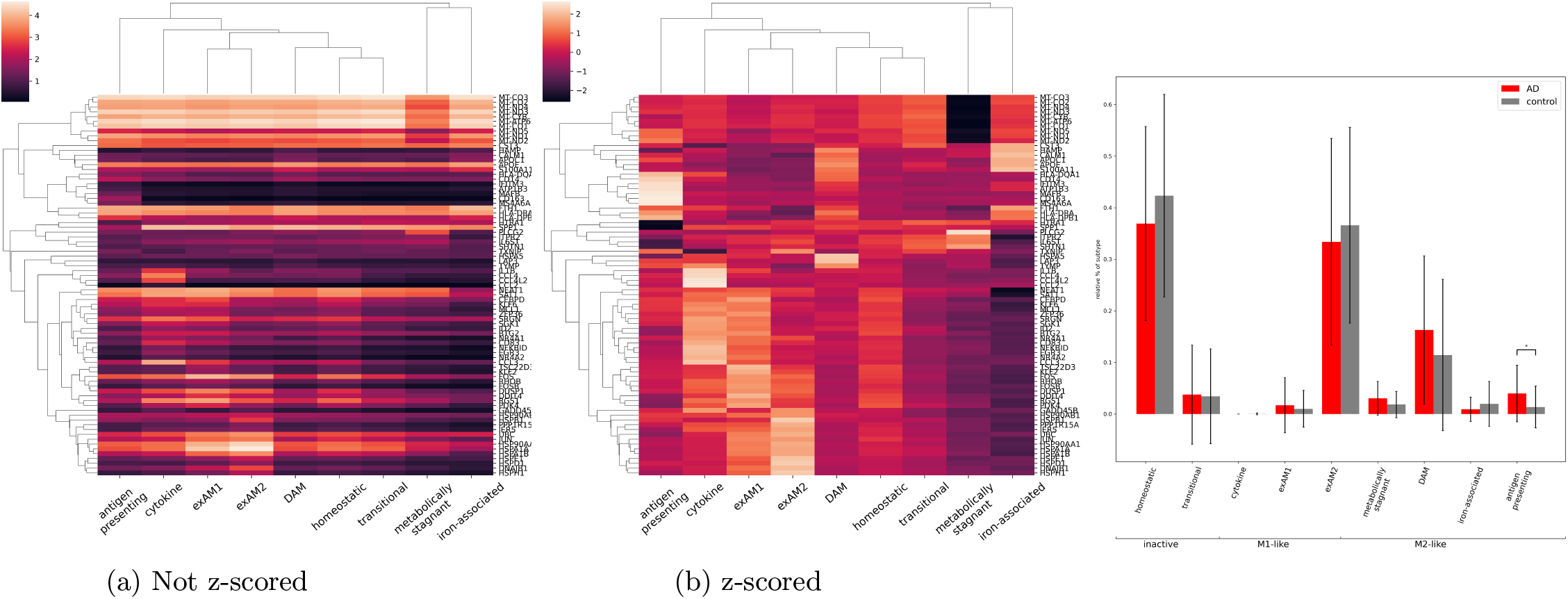
Clustermaps of log(CPTT+1) values of our signature genes without (a) and with (b) z-scoring within each cluster. Row and column dendrograms were created using the z-scored data. These genes are differentially expressed in the given cluster, are expressed in at least 50% of cells, and are among the genes with the largest variance in their SHAP values. (c) Subtype percentages between Alzheimer’s patients (AD) and controls on bulk, FACS-sorted microglia [40] after deconvolution using sampled signature matrices. Heights denote average percentage while error bars denote standard deviations.

The combination of SHAP and differential expression allows us to deduce the possible function of these clusters. Our approach yielded a total of 9 subtypes, many of which have similar gene expression patterns and characteristics of previously reported microglial subtypes. As previously mentioned, because of the variability in parameters used and experimental approaches employed, the gene lists for each subtype can vary across studies. That said, there are similarities across studies for many of the identified subtypes. An additional challenge in characterizing these subtypes is that they are understandably described in an effort to elucidate their underlying function or biology within the brain and disease. As the gene lists can differ, so can the inferred function. We are subject to the same interpretational challenges, even with our approach being grounded in an explicitly articulated biophysical approach.

We named the 9 subtypes as follows: 1) homeostatic, 2) transitional, 3) metabolically stagnant, 4) cytokine, 5) exAM1, 6) exAM2, 7) DAM, 8) iron-associated, 9) antigen presenting. Here we will touch briefly on the transcriptional signatures of the cluster, while a more detailed discussion of each subtype’s possible function, their relation to each other, and how they might impact Alzheimer’s pathology can be found in the Discussion.

Our homeostatic cluster has, as expected, a weak differential expression profile, and the largest fraction of high *I*_*c*_ cells. In fact, this cluster contains no unique differentially expressed genes. Most of the important SHAP genes exhibit clear transcriptional patterns in other clusters, suggesting their low/high SHAP values are due to high/low values in other clusters. A similar pattern in the transitional cluster, may, given the absence of of high *I*_*c*_ cells, indicate that the transition is being completed, and cells are heading to their final state.

The metabolically stagnant cluster is an especially interesting case, as we will discuss below. Among those that are up regulated is PLCG2 which has been implicated in AD with specific variants resulting in a protective, hyper-morphic phenotype [34, 35, 36]. ITPR2, another gene up-regulated in this cluster with a positive SHAP value, is a downstream effector of PLCG2 that stimulates the release of Ca2+ from the endoplasmic reticulum. The strongest signature of this cluster is the down-regulation of nearly all the genes of the mitochondrial genome, which have positive SHAP for low values.

The cytokine cluster shows strong signs of chemokine release, as seen through significant upregulation of CCL3, CCL4, CCL4L2, IL1*β*, NFKBID, and CD83 transcripts. CD83 is an marker for microglia activation and reduced inflammation [37]. Work by our collaborators have identified expression of CD83 to be found in patients in which CMV was detected [38]. This cluster, in which chemokine genes also display large SHAP values, was one of the most distinct and long lived cluster based on its variance along the third diffusion coordinate. Interestingly, almost half of the cells in this cluster are from a single sample from a patient in the Patel data set who had cancer, and was collected from the margins of a resected brain tumor.

Marsh et al. identified a gene expression signature in microglia that was dependent on the method of tissue dissociation used, termed the ex vivo activated microglia (exAM1 and exAM2) signature [39]. We observed two substates with a high overlap of these signatures, with differentially expressed genes and high SHAP genes being almost identical. For instance, both of these show upregulation of heatshock proteins HSPA1A, HSPA1B, HSP90AA1, HSP90AB1, HSPD1, HSPE1, HSPH1, HSPB1,and DNAJB1. Additional differentially expressed genes are DUSP1, UBC, the transcription factor JUN, and DNA damage inducible transcript DDIT4. These also appear in our SHAP analysis, indicating that high expression of these genes has a positive impact on classification into this cluster. This cluster has an over representation of cells from the Patel data set, which used enzymatic dissociation [2] as opposed to Olah set which used mechanical dissociation [1]. As such, we identify this cluster with the *ex-vivo* activated microglia (exAM) identified in [39]. We also performed differential expression analysis between these two clusters to try and uncover differences between them. The results suggest that exAM2 expresses higher level of heatshock proteins, while exAM1 expresses higher levels of various transcription factors (KLF6, KLF2, FOS, JUNB, JUND, FOSB) as well as cytokines (CCL2, CCL3).

The disease associated microglia (DAM) cluster is named as such due to the differential expression and large SHAP values for a number of the canonical DAM genes, including APOE, APOC1, CST3, and SPP1. And while these genes are not unique to this cluster, we observe high expression of these genes, as well as CD9 in a subset of cells in this cluster. CD9 did not meet the 50% threshold to be considered a signature gene, but is uniquely expressed in a subset of the cells in this cluster.

The iron-associated cluster in figure 1 has most of its differentially expressed genes downregulated and have positive SHAP for low values of the features. Specifically, we see downregulation of many cytokines and microglia activation markers, such as CD83, CCL4 and NEAT1. Among the up regulated genes are APOE, APOC1 and CST3, which have been studied in the context of AD before and have been associated to the disease associated microglia (DAM) phenotype. However, the other canonical DAM genes are down regulated in this cluster, including SPP1, hinting at a similar but distinct function from DAM. MARCKS, HAMP and FTH1 are also upregulated in this cluster, the latter two implying increased iron accumulation and metabolism in this subtype.

Our antigen-presenting cluster shows macrophage specific markers such as CD163, LYZ, and VIM, are shown to be highly upregulated in this cluster, although the percent of cells in which the latter two are expressed do not meet the 50% threshold. Of the genes which do meet the 50% cuttoff, the highest upregulated was MS4A6A, an AD-GWAS gene, followed by CD14. We also see overexpression and high SHAP for several the MHC class II genes (HLA-DQA1,HLA-DQB1, HLA-DPB1 and HLA-DRA), which present antigens to T-cells during immune response, as well as interferon induced transmembrane protein 3 (IFITM3).

### Comparison to previous results

One difficulty in quantitative comparison across different analyses is the methods and software packages used to determine the unsupervised clustering and marker genes. While we’ve attempted to keep our preprocessing in line with their reported methods, differences in analysis will lead to differences in results. For instance, in their initial analysis of this combined dataset the authors of [2] identified 12 unique subtypes based off of Leiden clustering on a UMAP-type nearest neighbor graph, and differential analysis performed in Seurat [20]. The strength of the FP diffusion maps is in their ability to i) accurately represent the dynamics of subtype differentiation [13], and ii) to identify robust long lived clusters by looking at a cluster’s variance along the first couple of diffusion coordinates. Based on our results in figure 1, we see that these robust clusters are the cytokine, antigen presenting, metabolically stagnant and anti-inflammatory clusters. The DAM, homeostatic, transitional and exAM clusters are more susceptible to noise and therefore more difficult to draw the boundaries between. When comparing the clusters from both analyses as in table 1, we see a substantial overlap between many of our clusters and Patel’s clusters. A lot of these agree, such as the overlap between our cytokine cluster and their leukocyte recruiting cluster, our exAM clusters and their stress induced clusters, our antigen presenting and their immunoreactive cluster, our iron-associated and their iron-accumulating clusters, and our DAM clusters. Interestingly, our metabolically stagnant cluster had the highest overlap with their homeostatic cluster. This may be due to our interpretation of the implication of our genes, or to our unique transcriptional signature which associates this cluster with increased IP3 production and calcium overload in the mitochondria, which is a nonstandard pathway to induce senescence. Interestingly, the highest differentially expressed gene for this cluster is PLCG2, which has an average log2 foldchange three times larger than the next largest foldchange. Additionally, there are several genes associated to A*β*, which does suggest increased IP3 production, possibly due to plaque uptake by TREM2.

**Table 1:** Percentage of cells in Patel clusters [2] found in our clusters. Rows add to 1. We find high overlap in our robust, long lived clusters and smaller overlap in others. Bold-faced type denotes high overlap.

|  | Cytokine | Homeo | Trans | Met. Stag. | exAM1 | DAM | exAM2 | Antiinfl. | Antigen pres. |
| --- | --- | --- | --- | --- | --- | --- | --- | --- | --- |
| Unknown | 0.1151 | 0.176 | 0.2401 | 0.1532 | 0.0254 | 0.1647 | 0.0468 | 0.0709 | 0.008 |
| FOS+ | 0.1981 | 0.2582 | 0.0574 | 0.0391 | 0.3603 | 0.0418 | 0.0385 | 0.0036 | 0.0029 |
| Unknown | 0.0579 | 0.1689 | 0.347 | 0.2441 | 0.0347 | 0.0479 | 0.0608 | 0.0328 | 0.0059 |
| Leuk. recruit. | **0.8393** | 0.0385 | 0.0048 | 0.0128 | 0.0774 | 0.0189 | 0.0043 | 0.0009 | 0.003 |
| Stress ind. | 0.0773 | 0.0437 | 0.0562 | 0.0385 | 0.1511 | 0.0082 | **0.6229** | 0.0012 | 0.0009 |
| Immuno | 0.1551 | 0.0556 | 0.0358 | 0.043 | 0.013 | 0.1218 | 0.0126 | 0.0367 | **0.5263** |
| Homeo | 0.0046 | 0.0037 | 0.2268 | **0.7137** | 0.0009 | 0.0028 | 0.0143 | 0.0332 | 0.0 |
| Iron accum. | 0.0067 | 0.0385 | 0.1682 | 0.1481 | 0.0008 | 0.0335 | 0.005 | **0.5824** | 0.0167 |
| Macrophage | 0.0653 | 0.0503 | 0.1256 | 0.0503 | 0.0352 | 0.206 | 0.0 | 0.2111 | 0.2563 |
| DAM-like | 0.1782 | 0.0922 | 0.0707 | 0.0445 | 0.0123 | **0.4839** | 0.0292 | 0.063 | 0.0261 |
| FOS+/Stress | 0.0101 | 0.0101 | 0.0 | 0.0 | **0.5808** | 0.0 | 0.3939 | 0.0 | 0.0051 |
| Proliferating | 0.1325 | 0.0361 | 0.1446 | 0.0843 | 0.012 | 0.1205 | 0.0361 | 0.3855 | 0.0482 |

### Deconvolution of an independent bulk microglia show antigen presenting subtype enriched for AD patients

Our analysis so far has been performed on a mixed data set with fresh and postmortem tissues from patients with various pathologies. One issue in using our set to observe various subtype’s association to AD is that all of our AD samples were from postmortem tissue, while most of the controls are fresh. In order to address this, we perform deconvolution using CIBERSORTx on an independent, bulk FACS-sorted microglia data set published by Kosoy et. al. [40] using our signature genes as our cellular subtype signatures. As discussed above, these genes are differentially expressed in each cluster, expressed in at lead 50% of cells in that cluster, and must have large variance in their SHAP values for the cluster (figure 7) This data set has considerably more power to distinguish differences in subtype levels with 55 AD patients and 57 healthy controls, compared to the 17 total and 6 total from Olah and Patel, respectively. Our results are displayed in table 2 and figure 7. We found AD patients to be most enriched in antigen presenting microglia. This result is especially interesting in consideration of the recent report suggesting recruitment of T-cells into the brain by microglia leads to increased tau and neurodegeneration [41]. Additionally, the p-value for the difference in DAM microglia barely missed the significance threshold, going in the expected direction (p-value = 0.06), as does the difference in the senescence subtype (p-value = 0.14).

**Table 2:** Subtype percentages between Alzheimer’s patients (AD) and controls on bulk, FACS-sorted microglia [40] after deconvolution using sampled signature matrices.

|  | homeo | trans | met. stag. | cytokine | exAM1 | exAM2 | DAM | iron | ant. pres. |
| --- | --- | --- | --- | --- | --- | --- | --- | --- | --- |
| mean % AD | 36.92 | 3.75 | 3.04 | 0.0 | 1.69 | 33.40 | 16.29 | 0.92 | 3.98 |
| mean % control | 42.36 | 3.42 | 1.84 | 0.02 | 1.00 | 36.61 | 11.42 | 1.98 | 1.35 |
| KS statistic | 0.14 | 0.05 | 0.23 | 0.04 | 0.08 | 0.14 | .26 | .12 | 0.34 |
| p value | 0.70 | 0.99 | 0.14 | 0.99 | 0.99 | 0.69 | 0.06 | 0.84 | **0.005** |

## Discussion

In this work we interrogated the dynamics of microglia substates with the purpose of (i) elucidating the dynamical structure of microglia subpopulations, (ii) determine which subpopulations exhibit clear dynamics separate from noise, (iii) determine genes associated to these clusters, and (iv) looked for subtype percentage differences by deconvolving an independent bulk microglia data set. To do this, we used the FP diffusion maps to find a low dimensional representation of the dynamics. We performed Leiden clustering with a resolution parameter of 1, and identified differentially expressed genes using standard single cell methods. We built a machine learning classifier to assign cells to their subtypes and interpreted the results using SHAP values. Once the signature genes have been found, we used them to construct a subtype signature matrix so that we could deconvolve an independent, bulk microglia data set. We found that AD patients have more of the antigen presenting microglia cells compared to controls.

In terms of gene signatures, our approach yielded subtypes and genes that are consistent with previously reported work. Four of the subtypes (antigen presenting, DAM, metabolically stagnant and iron-associated) share down regulated genes. Intersecting these down-regulated genes and performing gene set enrichment with enrichR identified the known microglial pathways typically ascribed to anti-inflammatory M2 microglia: TNF*α* and NF-*κ*B. While only the antigen presenting subtype was statistically different between AD and control, the DAM subtype narrowly missed significance. The next closest significant subtype was the metabolically stagnant, with a p-value of 0.14. Downregulation of NFKB is also known to occur with omega-3 fatty acids eicosapentaenoic acid (EPA) and docosahexaenoic acid (DHA). This is consistent with the idea that the microglial subtypes where we see downregulation of TNFa and NFKB are subtypes that phagocytose cell membranes containing EPA and/or DHA. Based on the gene expression profiles of these subtypes, they all appear to be involved in debris clearance, as has been established with M2 microglia [42].

The antigen presenting cluster in figure 1 acts the most “macrophage-like” and found to be most significantly increased in AD individuals. Both Olah and Patel annotate macrophages that made it past the initial FACS sorting procedures. To increase the resolution of our microglia classification, we have excluded them from our analysis from the start. If we were to include them, however, they form their own cluster which is in close proximity to this cluster in the diffusion maps. The highest upregulated gene in this was MS4A6A, whose over expression has been shown to increase with soluble TREM2 in the cerebral spinal fluid [43] which has also been shown to be associated with tau pathology [44]. Forthcoming work by our collaborators identifies a role for IFITM3 in phagocytosis (under review), consistent with our interpretation of this subtype as “M2-like”. Recent work has shown that T-cell infiltration into the brain due to tau pathology can lead to increased neurodegeneration, potentially connecting our finding of increased antigen presenting microglia in AD patients related to this observation [41]. These antigen presenting microglia may have a role in activating these T-cells into harmful phenotypes which lead to neuronal death.

The disease associated microglia (DAM) subtype barely missed statistically significance between AD and control, with nearly 50% more of these cells in AD patients. While many of the aforementioned genes associated to this cluster are not unique, we observe high expression of these genes, as well as CD9 in a subset of cells in this cluster. One shortcoming of the community detection algorithm (Leiden) we used to find these genes is the resolution limit [45]. The resolution limit is a lower bound on the relationship between the sum of weights in a given cluster and the sum of the weights connecting the cluster to the rest of the graph. Below such a threshold, it is possible (but not guaranteed) for the algorithm to not detect the cluster. Its entirely possible that this is the case for the DAM cells, leading to their grouping with cells from an adjacent subtype, thereby corrupting the transcriptional signature. Specifically, we observed that the cells with the highest expression of DAM marker genes were only a small subset of the total cluster, leading to some canonical DAM genes being excluded from this cluster’s signature set. Nevertheless, there is still a strong signal from several canonical DAM genes.

The iron-associated subtype shares many similarities with the DAM subtype, but the unique gene hits paint a picture of iron and lipid metabolism. For instance, upregulation of MARCKS indicates an increased response to lippopolysaccharides (LPS) [46], while saturated fatty acids have been shown to upregulate HAMP [47] and become incorporated into myelin sheaths [48], implying this cluster may be related to lipid processing. HAMP is also a regulator of iron in humans and is known as key regulator of iron metabolism [49, 50]. Additionally, the upregulation of FTH1 implies increased iron accumulation in this subtype. Aside from being an inflammation marker, NEAT1 knockout mouse models of cancer have shown decreased tumor formation and metastasis due to a “switching off” of the final step in glycolysis [51]. The downregulation of NEAT1 in this cluster could be due to a similar mechanism. These genes together suggest this subtype is associated to an increased uptake of lipids and iron, which may again be due to phagocytosis of neuronal debris, with the relatively higher iron coming from the larger number of mitochondria in the axonal regions of neurons [52, 53].

The metabolically stagnant cluster is especially interesting. Crucially, PLCG2 is an integral part of the TREM2 signaling pathway [54, 55] which is important for clearance of myelin debris and A*β* plaques. This is facilitated by PLCG2’s involvement in Ca2+ signaling pathways downstream of TREM2. Further downstream in this pathway is ITPR2, which releases Ca2+ from the endoplasmic reticulum into either the cytosol or cytsolic-organelles. The up regulation of these genes, together with the dramatic down regulation of the mitochondrial genes related to oxidative phosphorylation (OXPHOS) could be considered strong evidence for cellular senescence. Recent work has shown that increased contacts between ITPR2 and mitochondria induces premature senescence by pumping calcium from the endoplasmic reticulum into the mitochondria, leading to calcium accumulation in the mitochondria which contributes to cell death [56, 57, 58]. A similar downregulation of the genes related to OXPHOS was found in recent work on lipid accumulating subtypes in microglia [59, 60], although it remains unclear if this is due to a mitochondrial dysfunction, or a switch from OXPHOS to glycolysis, as discussed in [59]. However, this switch is known to occur in proinflammatory subtypes [61, 62] and as discussed before, this subtype is more reminiscent of the anti-inflammatory M2 classification due to the down regulation of both inflammatory cytokines and their receptors, as well as transcription factors known to activate inflammatory pathways. Gene set enrichment of the DEGs using enrichR identified another neurodegenerative disorder, Charcot-Marie-Tooth (CMT) disease, in which contacts between the lysosome and mitochondria are dysregulated, also impacting calcium signaling [63, 64]. Interestingly, in CMT, this disruption occurs in neurons, suggesting that a common signal may be influencing these dynamics in both neurons and microglia.

One possible explanation for why we observed so few cytokine microglia is that both the physical and chemical dissociation techniques altered the cytokine gene expression profile into either exAM1 or exAM2. Given the prominent role of M1 inflammatory microglia in neurodegenerative disorders, we would assume that while the two observed exAM signatures appear to be somewhat artifactual, they are likely to have been in some form of M1 inflammatory state prior to their dissociation. For this and other possible reasons, we would anticipate that exAM subtypes are likely biologically relevant, even if their signatures are at least partially attributable to the dissociation technique used.

Our main motivation for this work was the lack of rigorous methods available for subtype identification. We believe our approach here is a step towards rectifying this by using *physical* models and scales to inform how, and why of a typical clustering analysis. For single cell data in particular, clustering using Fokker-Planck diffusion maps is a natural way to do this, since the dynamical theory of cell development is now well established in practice, and the Fokker-Planck equation is a physically and mathematically sound way of analyzing the long time behavior of stochastic systems. We hope that readers of this work will use and expand on this idea so that unsupervised clustering of single cell data can be put on solid theoretical ground and yield reproducible results. Of course, information is always lost in any dimensional reduction scheme and no method is perfect, but incorporating prior knowledge of physics and biology to guide dimensional reduction and unsupervised learning can help preserve the most important information.

A shortcoming of this analysis is intrinsic to all analyses of this sort: treating single cells as members of a continuous vector space. As discussed above, this approximation primarily captures the dynamics associated to highly expressed genes when the relative number of cells with low UMI genes (< 10^3^) is small. However given the subtle nature of these microglia cells, a full treatment of all the stochastic effects will likely improve our clustering results. Recent work on stochastic modeling of single cell data [14, 15] has instead advocated for modeling single cell dynamics using chemical master equations, although incorporating the effects of gene-gene interactions is still an open question. FP diffusion maps offer a natural setting for studying the relationship between these “microscopic” stochastic models and their continuum limits, via the connection to the Fokker-Planck equation and other methods of statistical mechanics and large deviation theory. Additionally, one can extend this work to non-isotropic graph kernels using the same methods.

In consideration of these limitations, and the continued generation of larger datasets, we are confident that the biology involved in understanding microglial dynamics and their role in Alzheimer’s etiology will continue to improve, with increasingly reproducible results.

## Methods

### Preprocessing the data and identification of long-lived states

We followed the preprocessing pipeline laid out in [2] as best as we could. Raw count matrices from both data sets were downloaded from Synapse (syn28450881 and syn22314351), and processed using Seurat v4 [65]. We first kept cells with total UMIs between 500 and 46425, total number of features less that 8000, and with less than 10 percent of the transcripts mapped to the mitochondrial genome. Each unique sample’s cells were then normalized to *log*(*CPTT* + 1) and the highly variable genes were identified using variance stabilizing transformation [66]. Once each sample’s highly variable genes have been identified, we integrate them using Seurat’s FindIntegrationAnchors() and IntegrateData() functions laid out in [20].

We then create a nearest neighbor graph using the pynnDescent python package which implements the methods of using the Fokker-Planck kernel for diffusion maps 12 (see: Supplementary Information). This yields a Markov chain which approximates the backward Fokker-Planck operator, and allows us to represent our dynamical system in low dimensions. We then run the Leiden community detection algorithm [68] with a resolution parameter of 1 directly on this nearest neighbor graph to identify the sybtypes.

### Fokker-Planck diffusion maps

A detailed description of Fokker-Planck diffusion maps with illustrative examples can be found in [13]. Here we provide a brief description of the key results. The idea is to find a low dimensional representation for the dynamics associated to a stochastic differential equation of the form

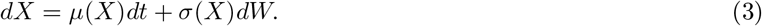

To do this, we exploit a theorem of Ting et. al [69] which derives an explicit expression for the drift and diffusion coefficients in terms of the local weights and bandwidths of a nearest-neighbor graph. In short, for an (isotropic) kernel of the form

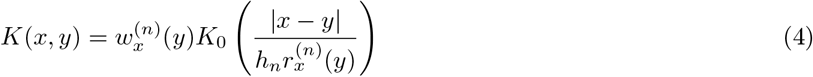

we have

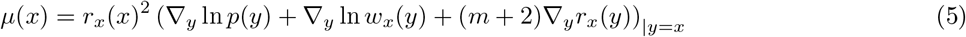

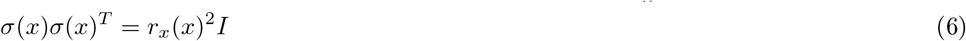

where *p*(*x*) is the sampling density.Next, we note that the diffusion approximations assumes that the system is in *local* equilbrium with respect to the potential. That is, that local density about each point takes the form of the equilibrium potential of the system described by eq. (1). We note that this method models the noise isotropic, i.e. it only depends on the absolute distance between points in transscriptional states. Incorporating anisotropic noise is possible, although it introduces several computational difficulties. For systems with multiplicative noise, and in the Stratonovich convention [70], this is

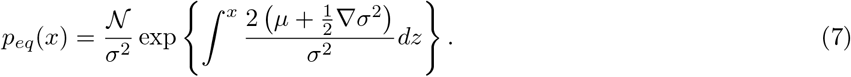

Since this is only expected to be true locally, we could approximate this using a local density estimate of the data data such as the kNN density as in [69], specifically we have

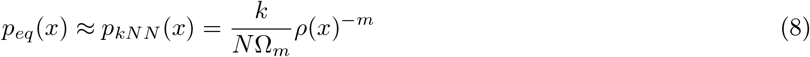

where Ω_*m*_ is the volume of the *m*-dimensional sphere, *N* is the sampling depth and *ρ*(*x*) is the distance to the *k*^*th*^ nearest neighbor. Using this, we can solve for the potential in terms of the noise and the distance to the *k*^*th*^ nearest neighbor. We find

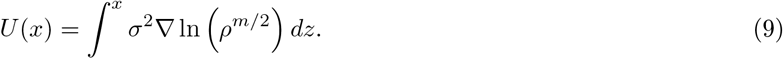

If we further estimate *σ*(*x*) using a linear function of the *k*^*th*^ nearest neighbor, we find

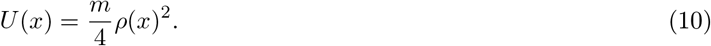

Thus, in order to approximate eq. (1) we need a normalization which gives

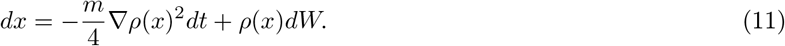

This motivates the following choice for our kernel

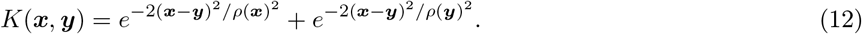

One advantage of using FP diffusion maps in this way is the identification of relaxation times associated to each diffusion coordinate. One way to solve the Fokker-Planck equation (more generally any linear PDE) is via the eigenfunction expansion, where the solution can be written as

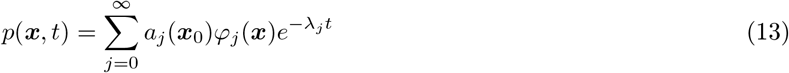

where *λ*_*j*_, *φ*_*j*_ are the *j*^*th*^ eigenvalue and eigenfunction, respectively, and *a*_*j*_(***x***_0_) is an unknown coefficient that depends on the initial conditions. In the presence of a spectral gap, i.e. when *λ*_*i*_ ≫ *λ*_*i*+1_ for some *i*, the dynamics only depends on *λ*_*j*<*i*_ since for times *t*≫ 1*/λ*_*i*+1_, the exponential factors for the other eigenvalues decay to zero. Above this spectral gap, the long lived (emergent) degrees of freedom are driving the dynamics, while the short lived (noisy) degrees of freedom equilibrate and become irrelevant. We can then isolate the long lived degrees of freedom from the noise, allowing us to identify robust substates. In the limit of small noise, these times are related to the size of the largest energy barrier needed to reach the global minimum [71]. Thus, the most prominent clusters in an FP diffusion map are those associated to the largest minima in our potential well, and are therefore the most robust to noise induced state-transitions.

### Calculating *I*_*c*_

To determine which cells are close to their transition point, we generalize the critical transition parameter first proposed in [27] into a field over each single cell. In it’s initial formulation, the proposed transition parameter for a set of cells (or states) *x*_1_, *x*_2_, ..*x*_*m*_ defined by a set of genes *g*_1_, …*g*_*n*_ at a given time *t* is

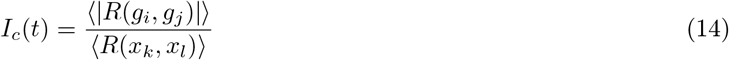

where *R* is the Pearson correlation coefficient and the average is performed over the off diagonal elements of the correlation matrix. Written another way, if ***X*** is our data matrix, where *X*_*ij*_ is the expression level of the *j*^*th*^ gene in the *i*^*th*^ cell, then we have

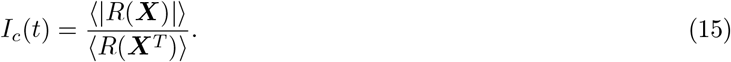

The intuition here, is that in when the system is in an attractor (the minimum of the potential), the denominator grows, since each cell has approximately the same gene expression profile, while the numerator shrinks. Alternatively, during a transition, certain genes become highly correlated as the cells move along a common trajectory from one attractor state to another. In this case, the numerator increase, while the denominator decreases. This intuition can be formalized using dynamical systems theory, which shows that eq. (14) diverges when the underlying dynamical system passes through a bifurcation point.

We generalize eq. (14) to a field over all cells by instead calculating the required correlation coefficients and averaging in a small neighborhood around each point *x*_*k*_. If we denote this neighborhood as Ω(*x*_*k*_), this gives

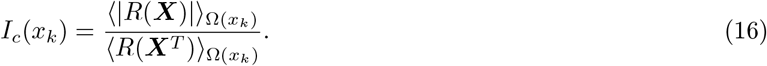

Given a pseudoordering of our cells *x*_*k*_, we can then study the time dependence of this quantity through time. After the cells have been appropriate ordered, we apply a non-uniform Savitzky-Golay filter [72], to smooth the resulting trajectory within each cluster, giving rise to figure 2.

### Differential expression and SHAP analysis

We use the default scanpy [32] method to perform our single cell differential expression. Once the genes have been found, we further filter to genes with an adjusted p value < .05, and genes which are expressed in at least 50% of the cells within that cluster.

To identify important genes for classification, we built a gradient boosted tree model using XGBoost [73] and interpreted the output using SHAP values [33]. We tuned our model using a randomized hyperparameter search with an 80-20 train test split. Our features were the diferentially expressed genes filtered according to the criteria in the previous paragraph, z-scored across all the genes.

### Deconvolution of bulk microglia data

We perform deconvolution on an independent bulk data using CIBERSORTx [74]. We built our signature matrix using the genes which both had large variance in their SHAP values and differentially expressed in a given cluster. We took the mean values of those genes across all the cells in that cluster. Default parameters were used when running the algorithm.

## Acknowledgements

This work was partially funded under US National Institute of Health Accelerating Medicine Partnerships-Alzheimer’s Disease (AMP-AD) consortium Grant No. 5U01AG046139 and by the Institute for Systems Biology.

## Data and code availability

Raw data used in this work is available through Synapse (syn28450881 and syn22314351). All scripts used in this work are available at: github.com/hadlocklab/FPdiffmaps/microglia.

